# Limitations of Variational Laplace-based Dynamic Causal Modelling for Multistable Cortical Circuits

**DOI:** 10.1101/2025.03.10.642327

**Authors:** Abdoreza Asadpour, Amin Azimi, KongFatt Wong-Lin

**Author notes:** These authors contributed equally to this work (Abdoreza Asadpour, Amin Azimi). Corresponding author: KongFatt Wong-Lin.

## Abstract

Dynamic causal modelling (DCM) is widely used to infer effective connectivity from neuroimaging data. However, its applicability to neural systems with complex, multistable dynamics remains uncertain—particularly when using the standard estimation approach based on variational Bayesian inference under the Laplace approximation. To investigate this limitation, we constructed biologically grounded cortical columnar neural mass models exhibiting three distinct multistable regimes: bistable fixed points associated with decision-making, coexisting oscillatory states through period-doubling bifurcations, and deterministic chaotic dynamics. These models were used to simulate local field potentials, which served as inputs to DCM. Bayesian model selection successfully identified the correct model architecture in all cases. However, Bayesian model averaging of the winning models failed to accurately estimate extrinsic connectivity parameters, leading to substantial discrepancies between the dynamics of the reconstructed and ground-truth systems. These results suggest that even when model selection is accurate, parameter estimation can break down under complex dynamics. Compared to previous applications of DCM to simpler neural systems, our study highlights significant limitations in its ability to capture the structure of multistable and globally nonlinear dynamics. We conclude that caution is warranted when applying variational Laplace-based DCM procedures to experimental paradigms involving bifurcations, chaotic trajectories, or other forms of dynamical complexity.

## 1. Introduction

Elucidating the functional and effective connectivity of healthy or dysfunctional brains is one of the aims in neuroscience (Friston 2011). In particular, network-based neuroscience offers opportunities to provide deeper understanding of the principles and mechanisms underlying complex brain function and cognition (Bassett and Sporns 2017). Functional and effective connectivity analyses are often conducted in human studies using non-invasive functional neuroimaging such as electroencephalography (EEG), magnetoencephalography (MEG), and functional magnetic resonance imaging (fMRI) (Friston 2011). In animal or certain drug treatment resistant clinical patient studies, functional and effective connectivity are applied to invasive intracranial/stereo EEG (i/sEEG) (e.g., Kobayashi et al. (2024)), local field potential (LFP) (e.g., Gallego-Carracedo et al. (2022); Meyer et al. (2018)) and multi-neuronal activity data (e.g., Tang et al. (2024); Trepka et al. (2022)).

Effective connectivity provides additional directed connectivity information than functional connectivity (Friston 2011) and is often state-space model dependent, for example, requiring the use of autoregressive models in Granger causality (Granger 1969), or nonlinear neural cortical (state-space) models in dynamic causal modelling (DCM) (Friston et al. 2003). In particular, DCM has been successfully applied to a variety of neuroimaging studies, not only in elucidating effective network connectivity but also inferring the underlying neural substrates for cognitive functions or dysfunctions (Huang and Nam 2020).

The models in DCM are based on the principle of canonical cortical columns (Mountcastle 1997). Within the DCM framework, effective connectivity is inferred through a combination of generative neural mass models, which simulate neuronal dynamics and observed neuroimaging data, and Bayesian statistical inference for model selection and parameter estimation (Friston et al. 2003; Penny et al. 2011). Specifically, standard DCM for event-related potentials (ERP), as originally introduced by David et al. (2006), employs variational Bayes inference under the Laplace approximation—known as variational Laplace—which estimates the posterior distributions of model parameters by maximising a quantity called negative variational free energy (Friston et al., 2007). While computationally efficient, the Laplace’s approximation inherently assumes a local, Gaussian distribution around the posterior mode (Friston et al. 2007). Consequently, it may fail to adequately capture posterior distributions that exhibit complex characteristics such as skewness, multimodality, or strong nonlinear dependencies, which are particularly likely to occur in neural systems with nonlinear and multistable dynamics (Chumbley et al. 2007). Thus, despite the widespread success of DCM, the reliability of its inference procedure—especially under complex neural dynamical states involving multiple coexisting stable states—remains unclear.

In this study, we assess how well standard time-domain ERP-based DCM can recover model architecture and connectivity strengths from simulated data reflecting multistable neural dynamics. This reflects a typical use-case scenario in which a non-expert user adopts the widely used inference procedure—Variational Laplace—without custom modifications (Zeidman et al. 2023). Such users typically do so because the true generative model is unknown, and ERP-based DCM is commonly assumed appropriate for paradigms with stimulus-locked inputs and expected evoked responses (Garrido et al. 2007). In contrast, for paradigms without well-defined inputs, such as resting-state activity, stochastic DCM is often more suitable (Li et al. 2011). Additionally, spectral DCM has been successfully applied to cases involving seizure dynamics (e.g., (Papadopoulou et al. 2015)) and cross-frequency coupling (Chen et al. 2008), where time-averaged spectral representations are more informative than event-related averages. Nevertheless, our goal here is not to compare DCM variants, but to highlight the limitations that may arise when a widely used time-domain ERP-based DCM is applied without modification to systems.

To simulate the data for this evaluation, we modified the generative neural mass models within the DCM framework to incorporate physiologically plausible dynamics with known ground truth. These adapted forward models generated synthetic neural data exhibiting three characteristic types of multistability: (i) bistable stable steady states (fixed points; i.e. choice attractors for decision-making (e.g. Roach et al. (2023a); Roxin and Ledberg (2008); Wong and Wang (2006)); (ii) two stable neural oscillatory frequencies (i.e. period-doubling condition) (Jia et al. 2012); and (iii) the extreme case where there is a finite continuum of multistable states i.e. deterministic chaos (Strogatz 2018). Importantly, these specific neural dynamics are investigated as they are not only physiologically plausible (Breakspear 2017) and can be generated from cortical column models, but they are also not determined by local dynamics about some steady state.

Although our results show that DCM can correctly identify the underlying model architecture within the specified model space, we also observed that the estimated connectivity strengths of the winning models often deviated from those of the ground-truth generative models. This discrepancy highlights the limitations of the variational Laplace-based DCM inference procedure—particularly under complex, multistable dynamical conditions. Our findings aim to promote a more informed application of ERP-based DCM and suggest that care should be taken when interpreting parameter estimates in systems with rich nonlinear dynamics.

## 2. Methods

### 2.1. Cortical columnar neural mass models

To evaluate DCM, we made use of local field potential (LFP) data generated from three different types of multistable neural mass models, which act as ground truths. In particular, all the models were based on cortical columnar structure conforming to DCM (Fig. 1).

**Fig. 1.**
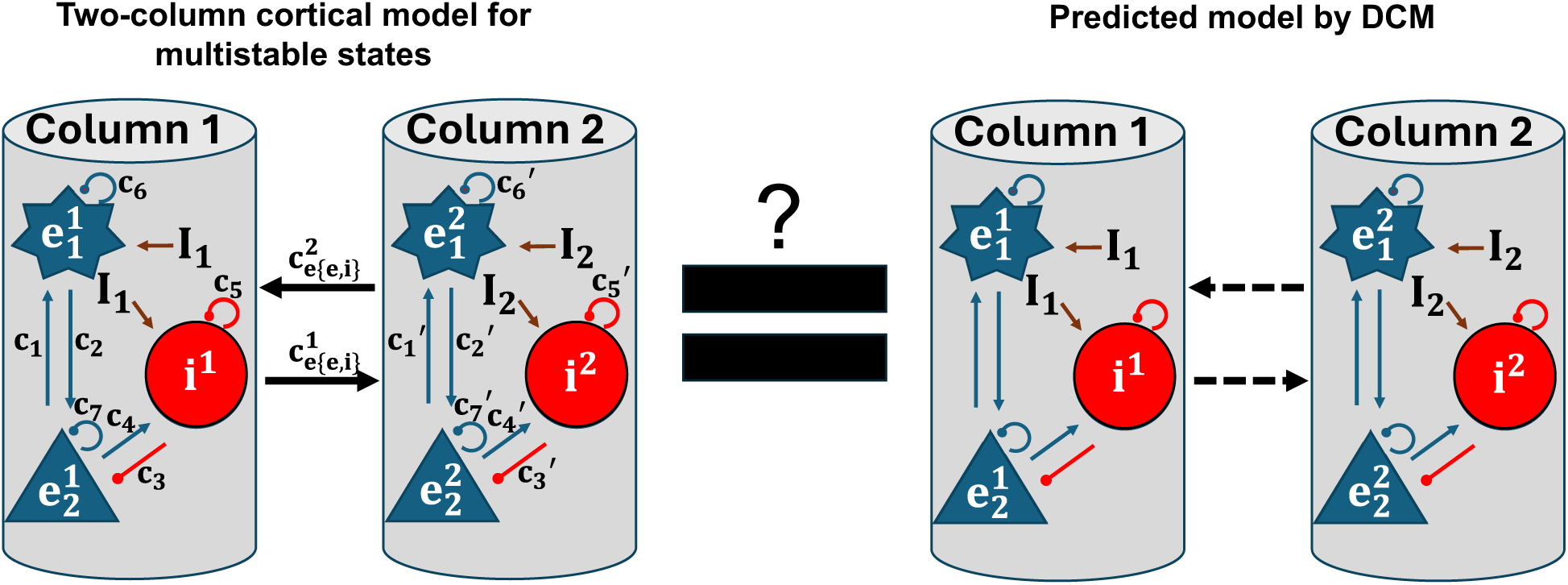
Schematic of DCM testing. To test whether the between-column (extrinsic) connections in multistable cortical circuit models (left) can be recovered by DCM-inferred models (right) in terms of model architecture identification and connectivity strengths based on the generated local field potential data from ground-truth models.

#### 2.1.1. Model with bistable fixed points that exhibits decision-making behaviour

The model with bistable fixed points for decision making comprised a two-column cortical neural mass model with two excitatory (*e*_1_ and *e*_2_) and one inhibitory (*i*) neural populations, lateral connectivity between columns and fully self-feedback, as described by Youssofzadeh et al. 2015.

We represented the stimulus input to both columns using a Gaussian bump function (Friston 2008):

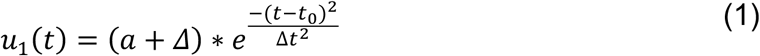

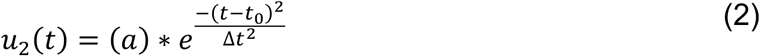

Here, *a* represents the amplitude at zero bias (*Δ*), with *t*_0_=0 s as the stimulus onset time and t=2 s as the stimulus offset time. The parameter *Δ*t (0.007 s) determines the width of the Gaussian bump.

Moreover, we define the evidence quality ε (analogous to motion coherence in random dot kinematogram) as follows:

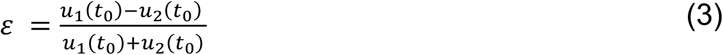

Noise is introduced to the excitatory (*e*_1_) and inhibitory (i) populations through the inputs *I_n_*, filtered via the fast synaptic activation of AMPA receptors. The dynamics of this process are described by the following equation (Wong and Wang 2006):

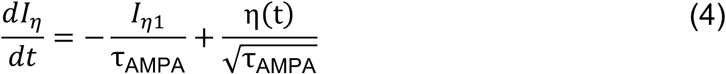

where *τ_AMPA_* is 5 ms, and *η*(*t*) is a white Gaussian noise with zero mean and a standard deviation of 0.02 nA. Then the total input current to each of the columns is:

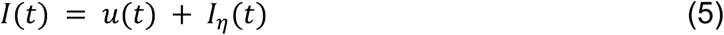

The network parameters were set to simulate two-alternative decision-making and working memory-based storage of the made decision via multistable dynamics as observed via postsynaptic potential (PSP) of *e*_1_ with respect to input current using XPPAUT, a widely used tool for simulating and analysing nonlinear dynamical systems (Ermentrout 2002). For DCM, we adjusted the input to the first column to be 50% greater than the input to the second column. We then categorised the trials into three types: (i) correct trials, where the output of the first column was significantly higher 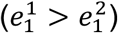 and reaching a decision threshold during stimulus; (ii) error trials, where the output of the second column was higher and reaching a decision threshold 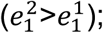; and (iii) ’non-decision’ trials, where the output difference between the two columns did not meet the decision threshold.

Within this model, the neural population’s mean firing rate function *S*(*v*), which resembles a sigmoidal curve, converts the mean post-synaptic potential *v* into the corresponding firing rate and is described by:

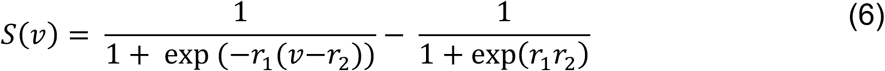

The parameters *r*_1_and *r*_2_ of the sigmoidal function influence its shape; increasing *r*_1_decreases the slope’s steepness, while increasing *r*_2_ moves the curve to the right. The mean firing rate, derived from the average membrane depolarization (or PSPs) through a static nonlinear sigmoidal function. Table 1 presents the priors and physiological meanings of the model parameters.

**Table 1.** Priors and the physiological interpretation of model parameters with fixed point bistability.

| Parameter | Physiological interpretation | Prior value (mean) |
| --- | --- | --- |
| $H_e, H_i$ [mV] | Excitatory and inhibitory magnitude in synaptic function | 3.25, 22 |
| $K_e = \frac{1}{\tau_e}, K_i = \frac{1}{\tau_i}$ [s<sup>-1</sup>] | Excitatory and inhibitory rate constant = 1/delay | 10, 200 |
| $c_1, c_2, c_3, c_4$ | Intrinsic coupling parameters for each column | 0.1, 0.1, 15, 3 |
| $c_5, c_6, c_7$ | Intrinsic coupling parameters in feedback loops for each column | 3, 0.1, 30 |
| $c_{ee}^1, c_{ei}^1, c_{ee}^2, c_{ei}^2$ | Extrinsic coupling parameters (lateral connections) | 0.6, 19, 0.6, 19 |

Subscript 'e' ('i'): excitatory (inhibitory)

According to schematic Fig. 2A, there is a lateral connection from neuron *e*_"_ in column one to neuron *e*_1_ in column two and vice versa. Moreover, there is a lateral connection from neuron *e*_2_ in column one to neuron *i*_1_ in column two and vice versa.

**Fig. 2.**
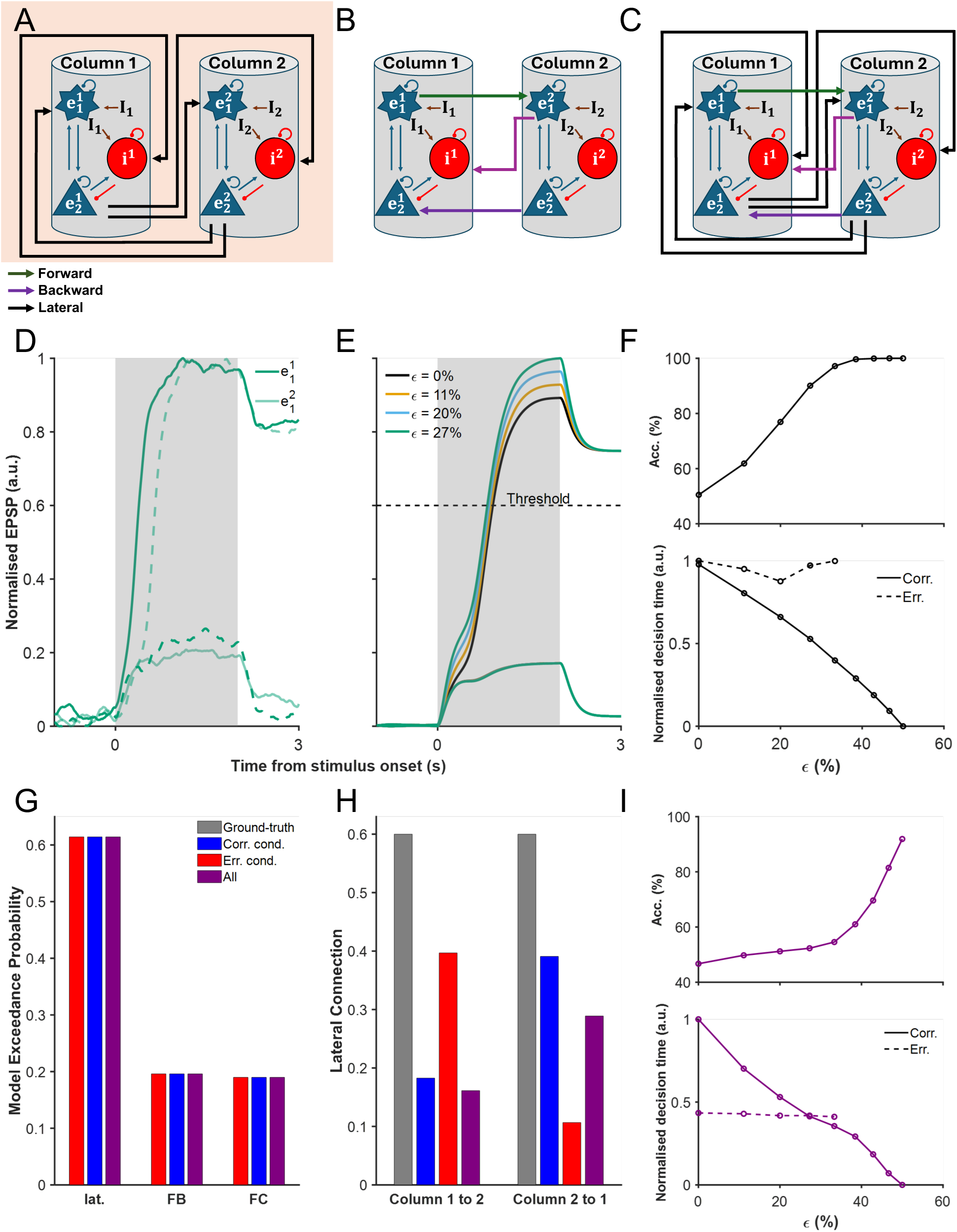
Evaluation of DCM’s ability to recover multistable cortical circuit dynamics. (A) Illustration of the ground-truth model, which incorporates only lateral (lat.) excitatory and inhibitory connections between cortical columns. (B) Alternative model featuring only forward-backward (FB) connections. (C) Fully connected (FC) model that includes both lat. and FB connections. (D) Simulated neural activity timecourse of *e*_1_ from the laterally connected model (A), with the output for column 1 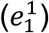 and column 2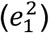 plotted separately. Solid (dashed) lines: correct (error) trials. (E) Neural activity timecourse across different input strengths (*ε*), with a decision threshold demarcating decision time for correct trials. (F) Psychophysical performance of the ground-truth model, where accuracy (Acc.; top) increases with input strength, while normalised decision time (bottom) decreases. (G) Bayesian model selection (BMS) results, illustrating the model exceedance probabilities across correct (Corr.), error (Err.), and combined (All) trial conditions. (H) The estimated lateral connection strengths inferred by DCM. (I) Reconstruction of accuracy and normalised decision time based on the recovered connection strengths of all-trial condition from (H).

Thus, a system of differential equations describing the two-column cortical model, according to Fig. 2, can be defined in the following.

For cortical column 1 (the superscripts in the equations below indicate the column number):

Post-synaptic potential for excitatory population 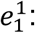

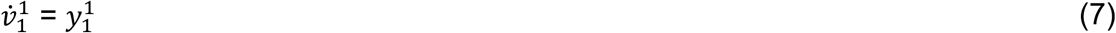

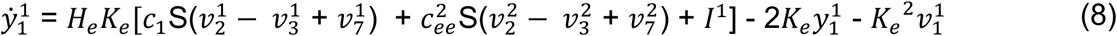

Post-synaptic potential for excitatory population 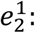

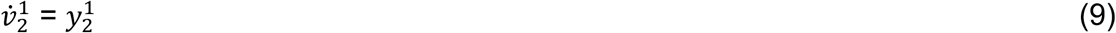

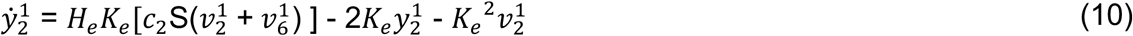

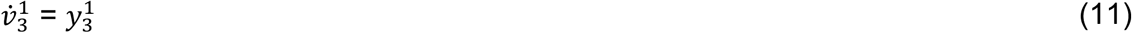

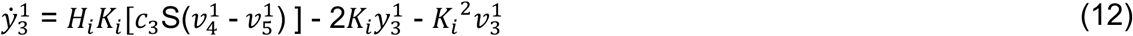

Post-synaptic potential for inhibitory population *i*^1^:

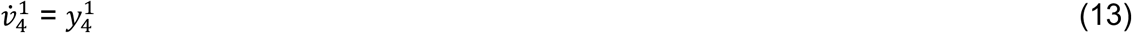

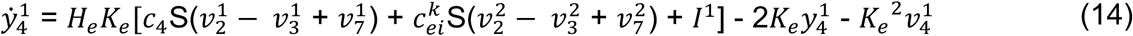

Post-synaptic potential due to self-inhibition for inhibitory population *i*^1^:

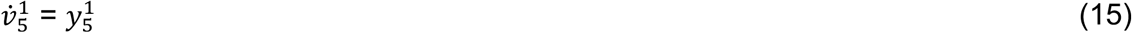

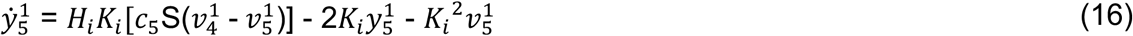

Post-synaptic potential due to self-excitation for excitatory population 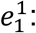

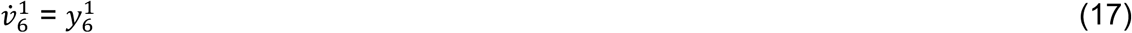

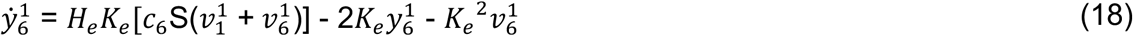

Post-synaptic potential due to self-excitation for excitatory population 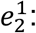

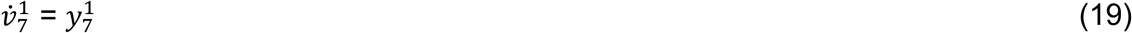

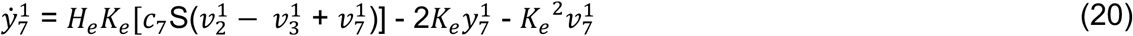

The LFP for column 1 is defined as (Mazzoni et al. 2015):

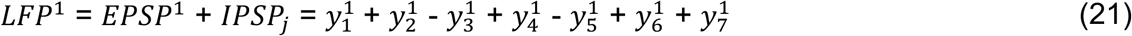

For the equations describing column 2, it is necessary to swap the superscripts in the corresponding equations for column 1 (Equations 7–21), changing superscript 1 to 2 and 2 to 1 accordingly.

#### 2.1.2. Models with period doubling and deterministic chaos

During period-doubling and chaotic regimes, we used three network architectures for DCM estimation to reveal distinct modes of cortical connectivity, such as decision-making bistability (Fig. 3A-C). To model the ground truth, we employed the model proposed by Ghorbani et al. (2012) to simulate the cortical column during period-doubling and chaotic regimes (Fig. 3A). However, due to the limitations of DCM in estimating synapses with temporal dynamics in the SPM12 software package (Penny et al. 2011), the adaptation for connections from excitatory-to-excitatory neurons is omitted.

**Fig. 3.**
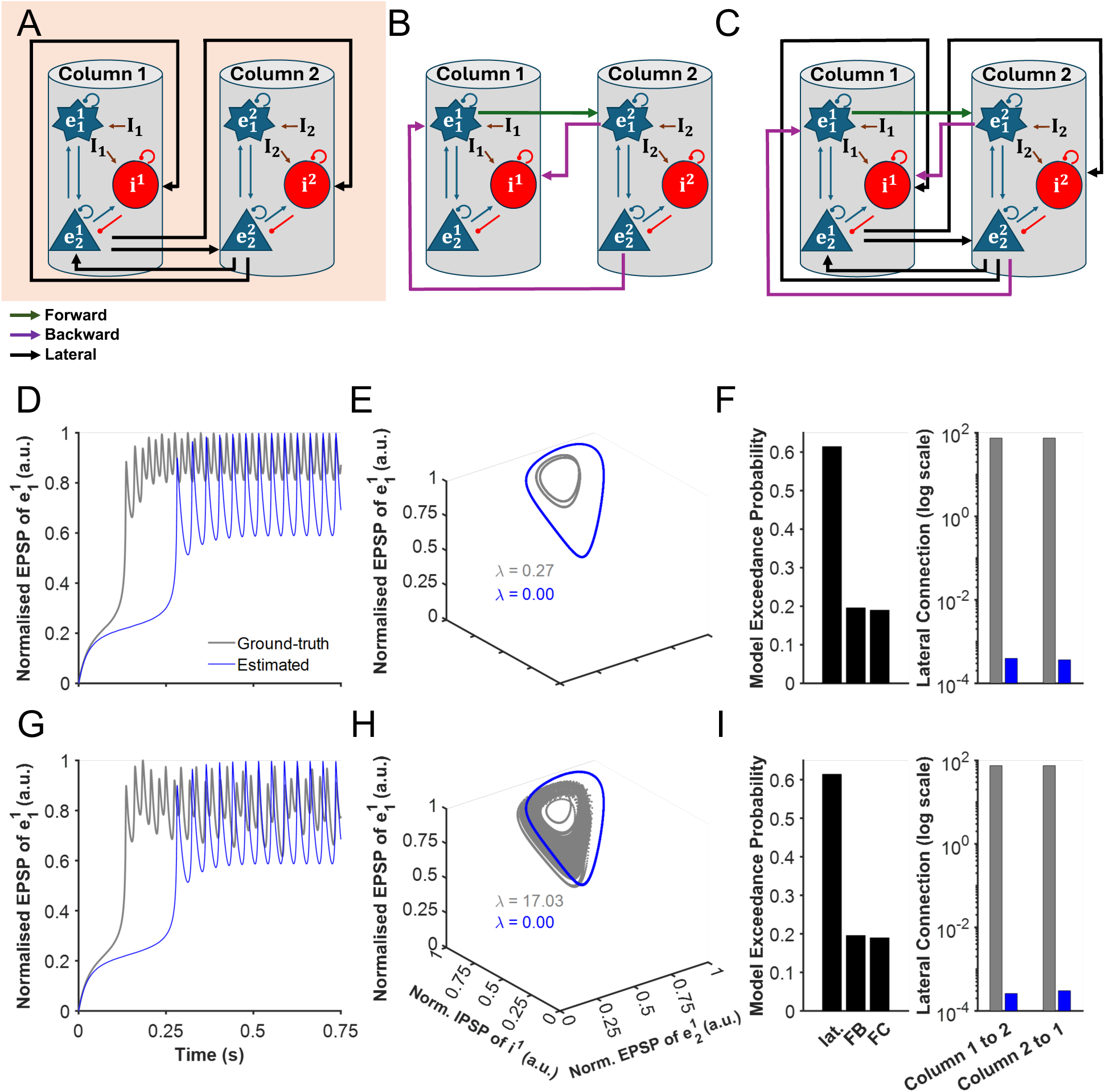
DCM estimation of period-doubling and chaotic dynamics. (A–C) Same as Fig. 2A-C. (D) Simulated neural activity of *e*_1_ in column 1 for the period-doubling regime, comparing the ground-truth model with the DCM-estimated model. (E) Phase-space trajectories for ground-truth and DCM-estimated (F) BMS results for period-doubling (left), alongside the recovered lateral connection strengths for ground-truth and DCM-estimated (right). G–I: Mirror (D–F) but for chaotic dynamics.

In this model, each neural population averaged membrane potential *V_m_* follows standard rate equation, where m denotes either excitatory (e) or inhibitory (i) neural population:

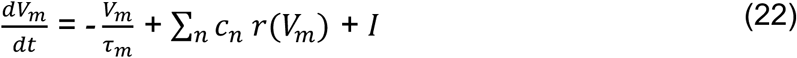

In this equation, the first term on the right-hand side describes the relaxation of the membrane potential to equilibrium. The membrane time constants *τ_e_* (20 ms for excitatory neurons) and *τ_i_* (10 ms for inhibitory neurons) represent the rate at which the membrane potential returns to its resting state after being depolarised or hyperpolarised (Bal and Destexhe 2009). The second term represents the changes in membrane potential due to excitatory post-synaptic potentials (EPSPs) and inhibitory post-synaptic potentials (IPSPs), where the sum runs over all neural populations connected to neural population n. Here, *r*(*V_m_*) is the firing rate of neural population m, and *c_n_* represents the connectivity strength between neural populations.

A neural population’s firing rate is determined by its membrane potential, which follows a sigmoidal curve as modelled by the following equation:

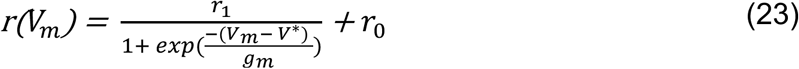

Here, *r*_1_ = 70 Hz is the maximum firing rate, *V*^∗^ is the firing threshold, and *g*_*m*_controls the steepness of the rate dependence on membrane potential. We assume that *g_e_* > *g*_i_, meaning that (e.g. fast-spiking) inhibitory neural populations have steeper firing rates in response to changes in membrane potential. In our simulations, we set *g_e_* = 5 mV and *g_i_* = 2 mV.

The model architecture for period-doubling and chaos comprises a two-column cortical neural mass model that includes two excitatory (*e*₁ and *e*₂) and one inhibitory (*i*) neural population, lateral connectivity between columns, and fully self-feedback connections. The nonlinear interaction between these two columns underpins period-doubling and chaos. Then the dynamics of the system are governed by three dynamical variables: the common membrane potential of excitatory neurons *V_e_*_1_ *and* *V_e_*_2_, the common membrane potential of inhibitory neurons *V*_1_. Their evolution is described by the following rate equations.

For cortical column j (the other column labelled as k):

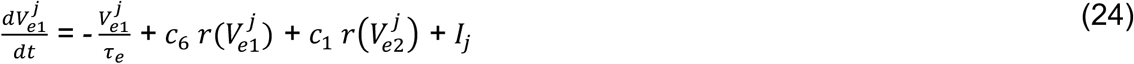

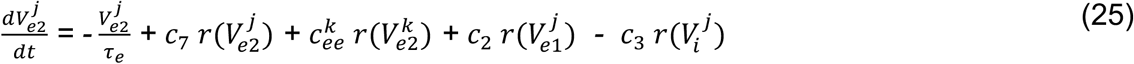

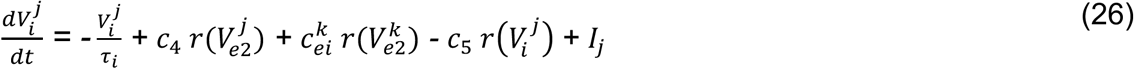

Tables 2 and 3 present the priors and physiological meanings of the model parameters during period-doubling and chaos.

**Table 2.** Priors and the physiological interpretation of model parameters with period-doubling.

| Parameter | Physiological interpretation | Prior value (mean) |
| --- | --- | --- |
| $c_1, c_2, c_3, c_4$ | Intrinsic coupling parameters for each column | 14.96, 14.96, 560, 760 |
| $c_5, c_6, c_7$ | Intrinsic coupling parameters in feedback loops for each column | 200, 14.96, 561.02 |
| $c_{ee}^1, c_{ei}^1, c_{ee}^2, c_{ei}^2$ | Extrinsic coupling parameters (lateral connections) | 74.80, 144, 74.80, 144 |

Subscript 'e' ('i'): excitatory (inhibitory)

**Table 3.** Priors and the physiological interpretation of model parameters with chaos.

| Parameter | Physiological interpretation | Prior value (mean) |
| --- | --- | --- |
| $c_1, c_2, c_3, c_4$ | Intrinsic coupling parameters for each column | 14.96, 14.96, 560, 760 |
| $c_5, c_6, c_7$ | Intrinsic coupling parameters in feedback loops for each column | 200, 14.96, 561.02 |
| $c_{ee}^1, c_{ei}^1, c_{ee}^2, c_{ei}^2$ | Extrinsic coupling parameters (lateral connections) | 74.80, 144, 74.80, 32 |

Subscript 'e' ('i'): excitatory (inhibitory)

In this model, since the definitions of EPSP and IPSP are not as straightforward as those in the decision-making model, we follow the methodology of Azimi et al. (2021) and define the LFP for each column as (- *V*_e2_).

### 2.2. DCM estimation of models

For DCM estimation, we assumed that the decision-making model conforms to an event-related potential (ERP) paradigm, as it involves stimulus-locked inputs and discrete response periods, matching the conditions under which ERP-based DCM is typically applied. Similarly, although the period-doubling and chaotic models exhibit more complex dynamics, they were also simulated over short epochs (1.5 seconds) with external inputs, which allowed us to treat them analogously as ERP-like responses for the purposes of model inversion.

Although our primary interest was in recovering the temporal dynamics of the period-doubling and chaotic regimes using a time-based DCM—motivated by the assumption of having defined external inputs and a short response window suitable for ERP-like modelling—we also implemented a CSD-based DCM estimation. This decision was guided by prior studies that successfully applied spectral DCM to seizure-like activity, which may similarly involve chaotic or period-doubling dynamics. We did not use stochastic DCM, as the presence of known inputs allowed us to inform the model structure directly and isolate the performance of the inversion procedure under multistable dynamics (see Supplementary Note 1 for methods).

To inform the generative model used by DCM, based on our alterations of intrinsic and extrinsic connections in the models, we modified the ERP-based DCM equations in the SPM software package (Penny et al. 2011) to match our revised cortical columnar formulas similar to Youssofzadeh et al. (2015). Specifically, for bistable decision-making model, we modified the state equations and prior moments for ERPs and leadfield parameters according to equations (1-(21 and Table 1. For period-doubling and deterministic chaos dynamics, we modified state equations and prior moments according to equations (22(26, Table 2, and Table 3. These adaptations ensured that the forward model reflected realistic and physiologically motivated connectivity structures consistent with each regime’s multistable dynamics.

Given current limitations in the DCM implementation in SPM, such as the lack of support for separate lateral connections to excitatory and inhibitory populations, we implemented a workaround in the chaos regime by scaling the inhibitory-to-excitatory connections in the toolbox to preserve the ground-truth ratio between excitatory and inhibitory lateral couplings.

For each simulation regime (bistable fixed point, period-doubling, and chaos), LFP trials were generated using only the corresponding ground-truth model with lateral connectivity. Based on the generated LFP, which estimates the neural population activities in the source (Penny et al. 2011), we first applied DCM based on three different possible models in model space. The first model was the ground-truth model with lateral connectivity between cortical columns (Fig. 2A and Fig. 3A) whereas the second model assumed the forward-backward connectivity between the columns (Fig. 2B and Fig. 3B) (with no modulatory connections). The third model is the fully connected model with both lateral and forward-backward connectivity (Fig. 2C and Fig. 3C). All DCM estimations were initialised using the default prior means provided by SPM software package, following standard practice in DCM-ERP studies (David et al. 2006). During model estimation, intrinsic parameters were fixed to their known ground-truth values across all simulations. Only extrinsic connectivity strengths were estimated using DCM, in order to specifically assess DCM’s ability to recover between-region (inter-column) connectivity under complex dynamical regimes.

### 2.3. Statistical analysis

#### 2.3.1. Bayesian model selection

After the DCM estimation in the model space separately for each trial, a Bayesian model selection (BMS) (Stephan et al. 2009; Wasserman 2000) identified the winning model using the random effects (RFX) procedure (Penny et al. 2011). We selected the RFX approach with the assumption of different performance mechanisms for different trials (Austin 2011). In particular, we treat each set of trials as coming from the same simulated "subject" across different "sessions", applying BMS using the RFX approach. This mimics standard group-level inference assumptions in DCM analyses and enables us to evaluate model performance under realistic conditions. This RFX method calculates the model probabilities which are described by a Dirichlet distribution and the optimised Dirichlet density using the observed data and ranking these model probabilities can compare the models at a group level (Stephan et al. 2009).

Although alternative group-level inference procedures such as parametric empirical Bayes (PEB) are increasingly used in recent DCM analyses (Zeidman et al. 2019a; Zeidman, et al. 2019b), we adopted RFX BMS here to remain consistent with the common usage among ERP-DCM applications—particularly those involving trial-wise model inversion, as is the case in decision paradigms. Moreover, our use of synthetic data allowed us to control variability, making RFX an appropriate and transparent choice for evaluating model identifiability. However, we implemented PEB wherever the RFX BMS could not recognise the ground truth as the winning model (see Supplementary Note 1). Model probabilities were summarised using exceedance probabilities, which quantify the belief that a given model is more likely than any other model in the space (Stephan et al. 2009). While exceedance probabilities have seen reduced emphasis in recent DCM literature in favour of log model evidence comparisons, we include them here to preserve comparability with earlier studies and standard ERP-based workflows. Nevertheless, we report model log evidences alongside for completeness.

Considering each model’s estimated posterior distribution given the observed data, one can utilise the Bayesian model averaging (BMA) strategy to calculate the average over all models by averaging the posterior distributions of models weighted by their posterior model probability (Trujillo-Barreto et al. 2004; Woolrich et al. 2009).

#### 2.3.2. Bayesian model averaging

To estimate the averaged extrinsic connectivity strengths for each condition, we performed BMA on the estimated DCMs of the winning model (Penny et al. 2011). This approach combines the parameter estimates from trials, weighted by the posterior probability of each model, to account for model uncertainty and provide robust averaged estimates.

While BMA remains a standard approach for summarising trial-wise DCM inferences, we acknowledge that more recent practices increasingly favour hierarchical inference through PEB, especially when evaluating effects across experimental conditions or subjects. In our work, which primarily focuses on trial-level inference, BMA is appropriate and consistent with typical ERP-DCM workflows used by non-expert users. However, we additionally employ PEB in cases where RFX BMS fails to identify the ground-truth model, in order to recover group-level effects and improve model discrimination.

#### 2.3.3. Reconstructed neural activity from DCM estimation

We simulated neural activity using estimated connectivity strengths of the winning model in BMS instead of the connectivity strengths in ground-truth models. For the bistable decision-making case, psychophysical metrics (decision accuracy and normalised decision time) were evaluated using simulations based on connectivity estimates obtained from correct and error trials combined. This combined-trial approach was eventually chosen because it has been demonstrated that decision-making models recover ground-truth parameters more effectively when using both correct and error trials, compared to when using either trial type alone (Asadpour and Wong-Lin 2024). For period-doubling and chaos regimes, we reconstructed and plotted the network activity timecourses and phase-space trajectories using the estimated BMA connections. For period-doubling and chaotic regimes, the Lyapunov exponents (Strogatz 2018) were computed.

### 2.4. Numerical method, and data and code availability

In numerical simulations for designing ground truth models before implementing DCM, the deterministic nonlinear differential equations for period-doubling and chaotic dynamics were solved using the Euler forward method, while stochastic differential equations for noisy bistable decision-making were solved using the Euler-Maruyama numerical method (Higham 2001). All simulations were performed in MATLAB R2023b software with a time step of 1 millisecond. Sufficient accuracy for these differential equations is achieved with Euler-based methods as long as the simulation time step used is sufficiently small, and it was verified that smaller time steps did not affect the results. For simulations using stochastic differential equations, the same set of pseudo-random seed was used when comparing between the generated data of the ground-truth models and corresponding DCM-estimated models, akin to Lenfesty et al. (2025). For DCM estimation, BMS, and BMA, computations were performed using MATLAB R2022a software via the Northern Ireland High-Performance Computing (NI-HPC) facility (https://www.ni-hpc.ac.uk).

## 3. Results

### 3.1. Cortical columnar model with multistable fixed points for decision making

We present a two-column cortical neural mass model that includes self-feedback (Youssofzadeh et al., 2015) and exhibits multistable fixed points for two-alternative decision-making (Roach et al. 2023b; Wong and Wang 2006).

We initially considered the input to two cortical columns to be random, assuming equal stimulus inputs in the form of Gaussian bumps (see Methods). In subsequent stages, we gradually increased the input Gaussian bump to one of the cortical column (column one) relative to the other cortical column (column two). A decision is considered made when the activity of one of the neuronal populations surpasses the threshold. By considering the model’s activity output, we observe three different types of simulated trials for various levels of inputs: correct trials, error trials, and non-decision trials (see Methods) (Fig. 2A and S1). Correct (error) trials occur when the cortical column that receives a higher (lower) stimulus input, that is column one (two), has its excitatory neural population reaches a prescribed decision activity threshold first, towards one of the two stable fixed points (a choice attractor (Roach et al. 2023b; Wong and Wang 2006)). Non-decision trials occur when none of the cortical columnar activities reach the decision threshold during stimulus presence.

The overall behaviour of the model recapitulated stereotypical psychophysics in two-choice reaction time experiments (Fig. 2D). Notably, the neural activity of the winning neuronal population exhibits a faster ramp-up with higher evidence quality (*ε*). Fig. 2E depicts the evolution of neural activity across different input strengths (ɛ), with the decision threshold clearly delineating the onset of the response for correct trials. Correspondingly, accuracy improves monotonically with increased evidence quality (Fig. 2F), while decision time decreases, with error decisions being slower than correct ones; typical speed-accuracy trade-off (Fig. 2F; Kaufman et al. (2015); Roitman and Shadlen (2002); Shadlen and Newsome (2001)).

To estimate DCM, we simulated the model across four distinct levels of evidence quality. For each level, we generated 200 correct trials and 200 error trials, resulting in a total of 400 trials per evidence quality level. This approach produced an aggregate of 1600 trials for comprehensive DCM estimation.

### 3.2. DCM identifies correct decision model architecture but not connectivity strengths

To evaluate the accuracy of DCM in identifying the underlying network architecture, we performed BMS across trials classified as correct, error, and all combined for the three different models in the model space with lateral connectivity, feedback connectivity, and full connectivity (Fig. 2A-C). The results show that variational Laplace-based DCM reliably identifies the ground-truth model as the most probable structure across all trial types (Fig. 2G). Notably, the exceedance probability of the model with extrinsic connections similar to ground-truth model remains significantly higher than that of the alternative models, suggesting that DCM effectively differentiates between model architectures in the given multistable cortical circuit. Additionally, mean Free Energy estimates confirm this preference, with the lateral model consistently yielding the highest evidence across trial types (Fig. S2A).

While BMS successfully selects the correct model architecture, the estimated extrinsic connection strengths obtained through BMA deviate from the ground truth (Fig. 2H). Despite correct model selection, the inferred lateral connection strengths differ from those in the simulated ground-truth model, suggesting that DCM struggles to recover precise network parameters. This discrepancy is particularly evident in trials where the winning model should maintain symmetric extrinsic connections, yet BMA estimates reveal noticeable asymmetries which in turn will introduce choice behavioural bias. Further, our results indicated that considering only correct or error trials alone lead to more asymmetry in connectivity strengths between columns than combining all trials.

To assess the impact of these estimation errors, we compared the psychophysical performance of the ground-truth model (Fig. 2F) with the performance reconstructed using DCM-estimated connection strengths for all trials by averaging over 1000 simulations for each evidence quality (Fig. 2I). The all-trials condition was used as it showed the least asymmetry (i.e. the best results) in connectivity between columns (Fig. 2H). When applying DCM-estimated connections from all trials, with combined correct and error trials, the reconstructed model’s decision accuracy deviated from that of the original model especially for intermediate evidence quality levels. Specifically, while the general trend was similar, the reconstructed accuracy curve differed in shape from the ground-truth, displaying a positive second derivative (concave upward) rather than the negative second derivative (concave downward) observed in the original data (compare Fig. 2I, top, with Fig. 2F, top). Furthermore, while the reconstructed normalised decision time for correct trials closely matched that of the ground-truth model, for error trials, the normalised decision time was substantially shorter across all evidence quality levels, indicating a consistent underestimation of decision times for error decisions (compare Fig. 2I, bottom, with Fig. 2F, bottom). These results highlight the inconsistencies introduced by DCM’s inversion procedure, potentially unable to make precise estimation to elucidate decision-making mechanisms.

### 3.3. Cortical columnar model with period-doubling dynamics

To further investigate DCM’s ability to infer multistable dynamics, we analysed a cortical model exhibiting period-doubling behaviour. The model space consisted of three candidate architectures: the ground-truth model with lateral connections, an alternative model with only feedback connections, and a fully connected model that includes both lateral and feedback connections (Fig. 3A-C). We trained DCMs using 400 simulated trials for each model in the model space.

In the ground-truth model, the simulated neural activity in column 1 (Fig. 3D, dark grey) exhibited characteristic period-doubling oscillations, where the system alternated between two stable oscillatory states. This behaviour was further reflected in the phase-space trajectory of the ground-truth model (Fig. 3E, dark grey), which showed a structured yet nonlinear pattern indicative of a system transitioning between distinct oscillatory regimes. This was further supported by its Lyapunov exponent value of 0.27.

### 3.4. DCM correctly identifies period-doubling model architecture but not connectivity strengths

BMS results (Fig. 3F) indicate that the ground-truth model is correctly identified as the most probable architecture. This is further supported by the Free Energy estimates from time-based ERP DCM, which also favour the lateral model over the other two architectures in the period-doubling regime (Fig. S2B, left). However, despite correctly selecting the model structure, DCM struggles to accurately recover the underlying connection strengths. Specifically, the inferred lateral connection strengths deviated substantially from those in the ground-truth model, suggesting that DCM did not precisely capture the intrinsic network dynamics governing period-doubling of the original cortical columnar model. Additionally, applying DCM-CSD to the period-doubling regime failed to recover the ground-truth model; both BMS, free energy, and PEB selected the forward-backward model instead (Fig. S3).

To assess the impact of these estimation errors, we simulated the network using the DCM-estimated connection strengths. The resulting neural activity timecourse (Fig. 3D, blue) deviated from the expected period-doubling behaviour. This suggests that the errors in the estimated connectivity disrupted the fine balance required for multistable oscillatory regimes. Similarly, the phase-space trajectory reconstructed using DCM-estimated connections (Fig. 3E, blue) lacked the well-defined structure of the ground-truth model, instead showing a trajectory indicative of a single stable oscillation, with Lyapunov exponent of zero.

These results highlight a critical limitation of DCM in estimating effective connectivity in networks exhibiting period-doubling. Similar to the decision-making case, while BMS can successfully identify the correct model architecture, inaccuracies in connection strength estimation compromise the ability of DCM to faithfully reconstruct the underlying oscillatory dynamics.

### 3.5. Cortical columnar model with chaotic dynamics

To investigate DCM’s ability to infer a continuum of stable dynamics, we analysed a cortical columnar model exhibiting deterministic chaos. The model space is the same as period-doubling (Fig. 3A-C). Unlike previous cases, the ground-truth model’s lateral connections to excitatory and inhibitory populations differ, which plays a critical role in generating the chaotic dynamics observed in this regime. As in previous case, we trained DCMs using 400 simulated trials for each model in the model space.

In the ground-truth model, the simulated neural activity timecourse in column 1 (Fig. 3G, dark grey) exhibited chaotic fluctuations characterised by aperiodic oscillations. This behaviour was further reflected in the phase-space trajectory (Fig. 3H, dark grey), which formed a finite, complex and non-repetitive structure, indicative of a high-dimensional chaotic attractor dynamics with a high Lyapunov exponent value of 17.03.

### 3.6. DCM correctly identifies chaotic model architecture but not connectivity strengths

BMS results (Fig. 3I) indicate that the ground-truth model was correctly identified as the most probable architecture. This is further supported by the Free Energy estimates from time-based ERP DCM, which also favour the lateral model over alternative architectures in the chaotic regime (Fig. S2B, right). However, for the chaotic regime, DCM-CSD again failed to consistently recover the ground-truth model. While Bayesian model selection and free energy comparisons favoured the forward-backward model, within-subject PEB correctly identified the lateral model as the most probable (Fig. S4), highlighting inconsistencies across inference methods. Similar to the previous cases, DCM struggled to accurately recover the underlying connection strengths. The inferred lateral connection strengths deviated substantially from those in the ground-truth model, suggesting that DCM did not effectively capture the nonlinear and highly sensitive interactions characteristic of chaotic dynamics.

To assess the impact of these estimation errors, we simulated the network using the DCM-estimated connection strengths. The resulting neural activity timecourse (Fig. 3G, blue) exhibited a qualitative shift, where chaotic fluctuations were replaced by oscillatory patterns that lacked the irregularity seen in the ground-truth model. This suggests that the errors in the estimated connectivity disrupted the delicate balance necessary for generating chaotic dynamics. Further, the phase-space trajectory reconstructed using DCM-estimated connections (Fig. 3H, blue) failed to reproduce the complex attractor structure of the ground-truth model, but instead exhibited a single stable oscillatory trajectory with a Lyapunov exponent value of zero. Hence, these results reinforce the limitations of DCM in estimating effective connectivity for multistable neural states.

## 4. Discussion

In this study, we evaluated the ability of DCM to infer effective connectivity in neural circuits exhibiting multistable dynamics. Using simulated local field potentials of cortical column models as ground truths, we investigated DCM’s performance in three cases of neural dynamics: bistable fixed points with decision-making, period doubling with two oscillatory frequencies co-existing, and deterministic chaos. Our results demonstrate that while BMS in DCM reliably identifies the correct model architecture across all cases, the standard DCM estimation procedure based on variational Bayes under the Laplace approximation struggles to accurately estimate extrinsic connection strengths. As only extrinsic parameters were estimated during model estimation, our findings directly reflect the reliability of DCM’s estimation of inter-regional connectivity under multistable dynamics, assuming correct intrinsic model specification. In this study, we purposely used local field potential source activity to avoid factors arising from volume conduction and delays (Sanchez Bornot et al. 2018), but DCM still struggles even with such case.

For the bistable decision-making case, we found that DCM-estimated connectivity strengths introduced systematic biases, affecting both decision accuracy and decision time. Similarly, in the period-doubling and chaotic cases, the estimated connectivity strengths failed to identify the network dynamics, leading to deviations from the network models’ activity timecourses, state-space trajectories and Lyapunov exponents from that of corresponding ground-truth models. Notably, combining both correct and error trials for BMA appeared to reduce asymmetry in the estimated extrinsic connectivity strengths between columns. We interpret this as a regularising effect, whereby pooling across decision outcomes provides a more representative sampling of the underlying dynamics and mitigates estimation bias introduced by unbalanced trial subsets. This is consistent with our previous findings using multivariate Granger causality for decision-making networks (Asadpour and Wong-Lin 2024).

A key limitation identified in our analyses concerns the performance of CSD-based DCM under complex dynamics. Specifically, under the period-doubling and chaotic regimes, model comparison using RFX BMS failed to identify the ground-truth architecture as the winning model. Notably, even when applying PEB to enhance group-level inference, the correct model could not be recovered in the period-doubling condition. This suggests that the assumptions underlying CSD DCM—such as time-invariant spectra and linearized dynamics—may be fundamentally misaligned with the nonlinearities and bifurcations characteristic of such regimes. These findings highlight a significant constraint on the applicability of CSD-based DCM to multistable neural systems. Accordingly, these results underscore the need for caution when applying CSD-based DCM to paradigms involving chaotic or period-doubling dynamics, such as seizure modelling, where standard assumptions may lead to systematic misinference.

Effective connectivity models are often used to infer directed interactions between brain regions (Friston 2011), and small errors in connectivity strength estimation can lead to incorrect conclusions about neural communication and information processing. Prior research has highlighted certain limitations of DCM, including its tendency to favour more complex models even when a simpler ground truth exists (Litvak et al. 2019), as well as difficulties in estimating synaptic gain which may result in Hopf bifurcations (Moran et al. 2013). Adding to previous findings, our results suggest that DCM’s estimation biases can lead to distorted behavioural outcomes. These findings suggest that DCM may not be fully reliable when applied to real-world neuroimaging data where multistability plays a central role, such as in processing decision-making (Wong and Wang 2006), working memory (Lundqvist et al. 2018), cognitive flexibility (Loh and Deco 2005), perceptual switching (Rankin et al. 2014), and pathological brain states like epilepsy (Kalitzin et al. 2019).

To further examine whether the observed DCM inaccuracies were intrinsic to the inference method or a property of the multistable regimes themselves, we conducted a control analysis in a single steady-state regime using the same thalamocortical equations but with balanced lateral couplings and increased self-excitation (c7 = 3) to suppress multistability (Fig. S5). This configuration represents the simplest model variant, with the fewest effective states and minimal intrinsic noise, and thus provides the most controlled case for testing DCM parameter recovery. Despite DCM underestimating the absolute magnitude of the lateral connections by several orders of magnitude, the predicted normalised EPSP closely followed the ground-truth trajectory, and the correct lateral architecture was selected by BMS. This finding indicates that, near a fixed point, DCM can accurately reproduce the observed temporal dynamics even with incorrect absolute parameter values, due to compensatory scaling among local excitatory and inhibitory gains and the model’s observation function – parameter degeneracy. The effect is further reinforced by SPM’s default data normalisation, which removes absolute amplitude information and allows multiple parameter sets to yield identical normalised trajectories. However, when the system enters a multistable regime, such compensation fails because the global nonlinear balance becomes essential for determining the emergent dynamics, leading to successful model-family identification but inaccurate parameter recovery. While this control demonstrates that DCM can match the observed dynamics under fixed-point conditions through parameter compensation, future studies should systematically investigate other types of single steady–state models and assess DCM’s ability to recover the ground truth connectivity more accurately.

An interesting auxiliary results we found was that DCM’s ability to estimate connectivity strengths in multistable cortical circuits was highly sensitive to the temporal resolution of the data (Fig. S6). Specifically, in our simulations, a sampling rate of 500 Hz—typical of EEG acquisition—proved insufficient for capturing the multistable dynamics during period-doubling and chaos. Increasing the rate to 10,000 Hz led to substantial improvements in DCM estimation. While we did not perform a systematic sweep across sampling frequencies, we acknowledge that such an analysis—e.g., plotting RMSE or estimation fidelity as a function of sampling rate—could be highly informative. This issue of insufficient sampling resolution may be particularly relevant in real EEG/MEG studies involving high-frequency neural dynamics, such as steady-state visually evoked potentials (SSVEPs), mismatch negativity (MMN), P300, and resting-state activity. Given that previous work has shown that DCM already faces challenges in capturing fast frequency components in neural signals (Friston et al. 2019), future studies using simulated or empirical data across a range of paradigms could systematically evaluate how sampling rate influences parameter recovery in DCM, particularly under regimes with fast or multistable dynamics. Addressing these challenges will require methodological refinements to DCM as well as the exploration of alternative or hybrid approaches that improve its capacity to capture multistable neural dynamics.

To improve DCM’s applicability to multistable neural systems, refinements in prior assumptions and model structure are necessary. One approach is utilising adaptive temporal filtering (Arnold et al. 1998) to better capture rapid neural state transitions. Moreover, the current reliance on the Laplace approximation assumes a Gaussian posterior, which may not adequately capture global nonlinear dependencies in complex systems (Chumbley et al. 2007). Using non-Gaussian priors or more flexible variational inference methods (e.g., Khan et al. (2013)) could enhance accuracy in estimating connection strengths. Non-Gaussian priors allow for greater flexibility in capturing skewed or multimodal posteriors that arise from nonlinear or multistable dynamics. For example, Chumbley et al. (2007) demonstrated using a Metropolis–Hastings approach that the true posterior distributions in DCM can exhibit significant non-Gaussian structure, particularly when priors are uninformative or the model is strongly nonlinear. In such cases, conventional Gaussian priors may over-constrain inference, whereas more expressive priors with bounded or skewed support may improve posterior recovery and reduce estimation bias. A stronger foundation for bifurcation-aware modelling could be achieved by incorporating dynamical priors that track bifurcations (e.g., Moran et al. (2011)) or machine-learning-based system identification (Wang et al. 2018). These improvements would help DCM better recover multistable dynamics. In addition, the use of multistart initialisation strategies—where the model inversion is repeated with different starting points—could mitigate the risk of convergence to poor local optima in the free energy landscape, thereby improving parameter recovery in systems with multimodal or non-convex posteriors.

To further understand the nature of estimation failures, we examined the evolution of free energy and its components—accuracy and complexity—across expectation maximisation iterations for representative failed cases in the period-doubling and chaotic regimes. This analysis demonstrates that although the models converge in terms of free energy, the internal dynamics of the estimation (particularly the accuracy term) evolve differently across regimes and architectures (Fig. S7). These findings reinforce that convergence alone does not imply successful recovery of the underlying dynamics, and that inspecting free energy components may help diagnose failure modes in DCM estimation.

In addition to variational Bayes-based inference, sampling-based approaches such as Markov Chain Monte Carlo (MCMC) (Hastings 1970; Metropolis et al. 1953), Hamiltonian Monte Carlo (HMC) (Duane et al. 1987), and Sequential Monte Carlo (SMC) (Gordon et al. 1993) provide asymptotically exact posterior estimates given sufficient samples. These methods, while computationally intensive, are especially suited for capturing complex, multimodal, or non-Gaussian posterior distributions that arise in multistable neural systems.

While refining DCM could enhance its ability to capture multistable neural dynamics, alternative approaches may offer complementary solutions. Granger causality (Granger 1969) provides an exploratory method but is limited to linear interactions and is prone to detecting spurious connections, making it less effective for capturing complex cortical dynamics (Asadpour and Wong-Lin 2024). Data-driven techniques such as recurrent neural networks (Downey et al. 2017) offer flexibility in identifying latent neural states but lack the interpretability of DCM’s structured parameter space. A promising direction is hybrid approaches that integrate DCM with data-driven inference, such as using variational autoencoders (Yang et al. 2023), dynamical systems based regression (Lenfesty et al. 2025), or Bayesian neural networks (Neil et al. 2007) to refine priors, improving estimation accuracy while preserving interpretability.

Taken together, we have evaluated the ability of DCM to infer effective connectivity in neural circuits exhibiting multistable dynamics. While BMS reliably identified the correct model architecture, the commonly used variational Laplace-based DCM inversion procedure struggled to accurately estimate extrinsic connection strengths, leading to deviations in simulated neural activity across decision-making, period-doubling, and chaotic regimes. These limitations suggest that while DCM remains a valuable tool for model selection, its application to multistable neural systems should be approached with caution. Future work should focus on refining DCM’s parameter estimation methods, incorporating bifurcation-aware modelling, and integrating data-driven approaches to improve its accuracy in capturing complex neural dynamics.

## CRediT authorship contribution statement

**Abdoreza Asadpour:** Conceptualization, Formal analysis, Validation, Visualization, Investigation, Methodology, Writing – original draft, Writing – review & editing. **Amin Azimi:** Formal analysis, Validation, Visualization, Methodology, Investigation, Writing – original draft, Writing – review & editing. **KongFatt Wong-Lin:** Conceptualization, Validation, Supervision, Project administration, Funding acquisition, Writing – original draft, Writing – review & editing.

## Code and data availability

The source code for our implementation and generated data are available and can be accessed at the following GitHub repository: https://github.com/asadpouretal/DCM_Multistability

This repository contains all the necessary scripts, models, and instructions to reproduce the results presented in this study, ensuring transparency and facilitating further research.

## Acknowledgements

A.A., AM.A. and K.W.-L. were supported by HSC R&D (STL/5540/19) and MRC (MC_PC_20020). We are grateful for access to the Tier 2 High-Performance Computing resources provided by the NI-HPC facility funded by the UK Engineering and Physical Sciences Research Council (EPSRC), Grant No. EP/T022175/1.

