## Supplemental Materials for "Limitations of Variational Laplace-based Dynamic Causal Modelling for Multistable Cortical Circuits"

[2]Current address: Sussex Neuroscience, School of Life Sciences, University of Sussex, Brighton, UK

[3]Current address: School of Psychology, Manchester Metropolitan University, Manchester, United Kingdom

Supplementary Material

This supplementary file contains additional details supporting the analyses and findings described in the main manuscript. Furthermore, Supplementary Note 1 describes the implementation of DCM-CSD for modelling cross-spectral densities from simulated period-doubling and chaotic dynamics.

- The supplementary figures provide further insight into simulation outcomes, model selection performance, and algorithmic behaviour under different regimes:Fig**. S1:** Example simulation of a non-decision trial within the bistable decision-making model.
- **Fig. S2:** Mean Free Energy comparisons across three model architectures (Lat, FB, FC) under ERP-based DCM for multistable, period-doubling, and chaotic regimes.
- **Fig. S3:** DCM-CSD results in the period-doubling regime, showing a failure to recover the ground-truth model.
- **Fig. S4:** DCM-CSD results in the chaotic regime, with only PEB recovering the correct model.
- **Fig. S5**: DCM can accurately reproduce the single steady-state output of the thalamocortical model despite strongly underestimated lateral connection strengths, owing to compensatory parameter scaling and the default data normalisation applied in SPM's ERP-DCM.
- **Fig. S6:** Demonstration of the impact of sampling frequency limitations (500 Hz) on DCM parameter estimation in the period-doubling regime, highlighting the inability of DCM predictions to accurately capture the original oscillatory dynamics despite algorithmic convergence.

- **Fig. S7:** Evolution of variational free energy, accuracy, and complexity across EM iterations for the three model architectures (Lat., FB, FC) under period-doubling (A) and chaotic (B) regimes in a sample trial. While all models converge, the decomposition profiles differ by regime, highlighting that convergence alone may not ensure accurate recovery of underlying dynamics.

### Supplementary Note 1. Implementation of DCM-CSD for Period-Doubling and Chaotic Regimes

This supplementary note outlines the computational steps used to implement Dynamic Causal Modelling (DCM) for cross-spectral density (CSD) analyses of simulated neural activity from cortical circuits exhibiting period-doubling and chaotic dynamics. All simulations and DCM estimations were conducted in MATLAB using a modified version of the SPM12 software package.

For both dynamical regimes, simulations were performed using cortical models with two columns, each consisting of interconnected excitatory and inhibitory neural populations. Simulated local field potentials (LFPs) were derived from these population activities. A high temporal resolution of 10,000 Hz was used, and multiple trials were generated with either low-amplitude stochastic inputs (for period-doubling) or sinusoidal waveforms modulated by noise (for chaos). The resulting time series were stored in structured data files for subsequent modelling.

DCM estimation was performed independently for each trial. First, cross-spectral densities were calculated from LFP data. Then, three competing DCMs were constructed, differing in their assumed extrinsic connectivity: (i) lateral only, (ii) forward-backward only, and (iii) fully connected models. These DCMs were configured with appropriate frequency windows, data ranges, and source structures, and were inverted using the standard CSD estimation routine available in SPM.

After model estimation, random-effects Bayesian model selection (BMS) was used to determine the most probable model architecture across trials. In addition, parametric empirical Bayes (PEB) was applied to perform second-level inference over model parameters. For each connectivity architecture, a group-level general linear model with a constant regressor was used to assess model evidence and review the best-fitting connectivity patterns.

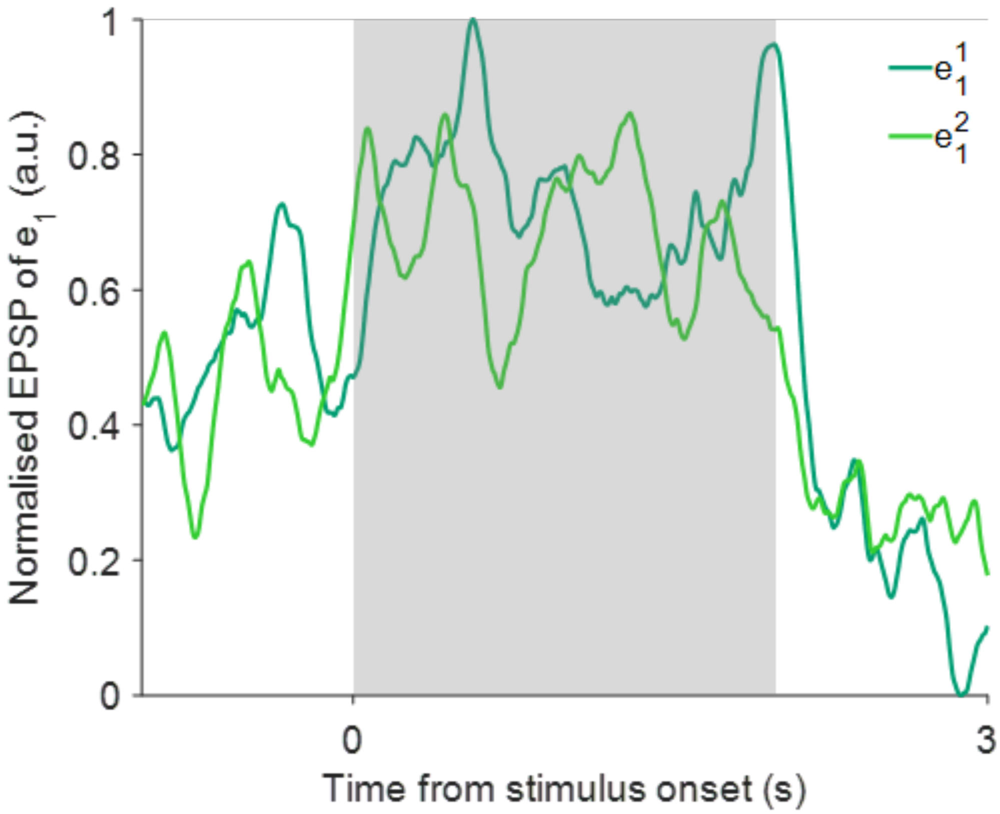

Figure S 1

**Fig. S1. Sample non-decision trial simulation of normalised EPSP (of neural populations $e_1$) from cortical columnar model with fixed point bistability.** Grey region: stimulus presence. $e_1^1$: normalised EPSP of neuron $e_1$ in column 1, $e_1^2$: normalised EPSP of neuron $e_1$ in column 2.

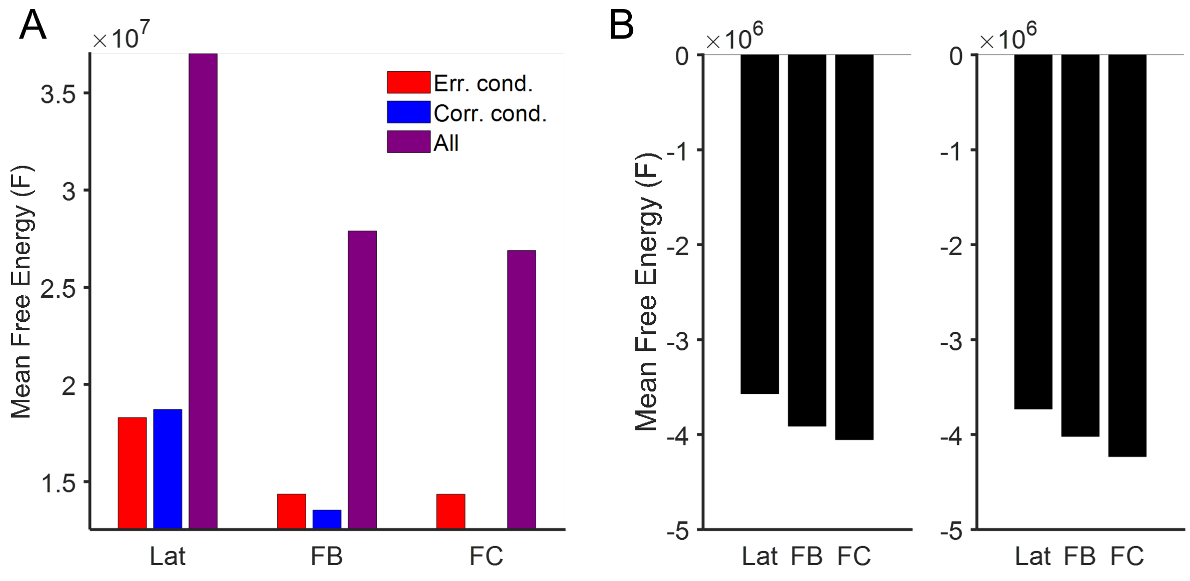


**Fig. S2. Mean Free Energy comparisons across models for time-based ERP DCM applied to simulated regimes.** This figure illustrates model evidence (Free Energy) for three network architectures—Lat (lateral connections only), FB (forward-backward connections), and FC (fully connected)—when using ERP-based DCM to model neural dynamics from simulated trials. **(A)** Free Energy estimates across error (red), correct (blue), and combined (purple) trials. The combined condition yields the highest overall Free Energy, with the lateral model (Lat) outperforming others across all conditions. **(B)** Mean Free Energy for each model in simulations of period-doubling (left) and chaotic (right) dynamics. In both cases, the lateral model (Lat) exhibits the highest evidence.

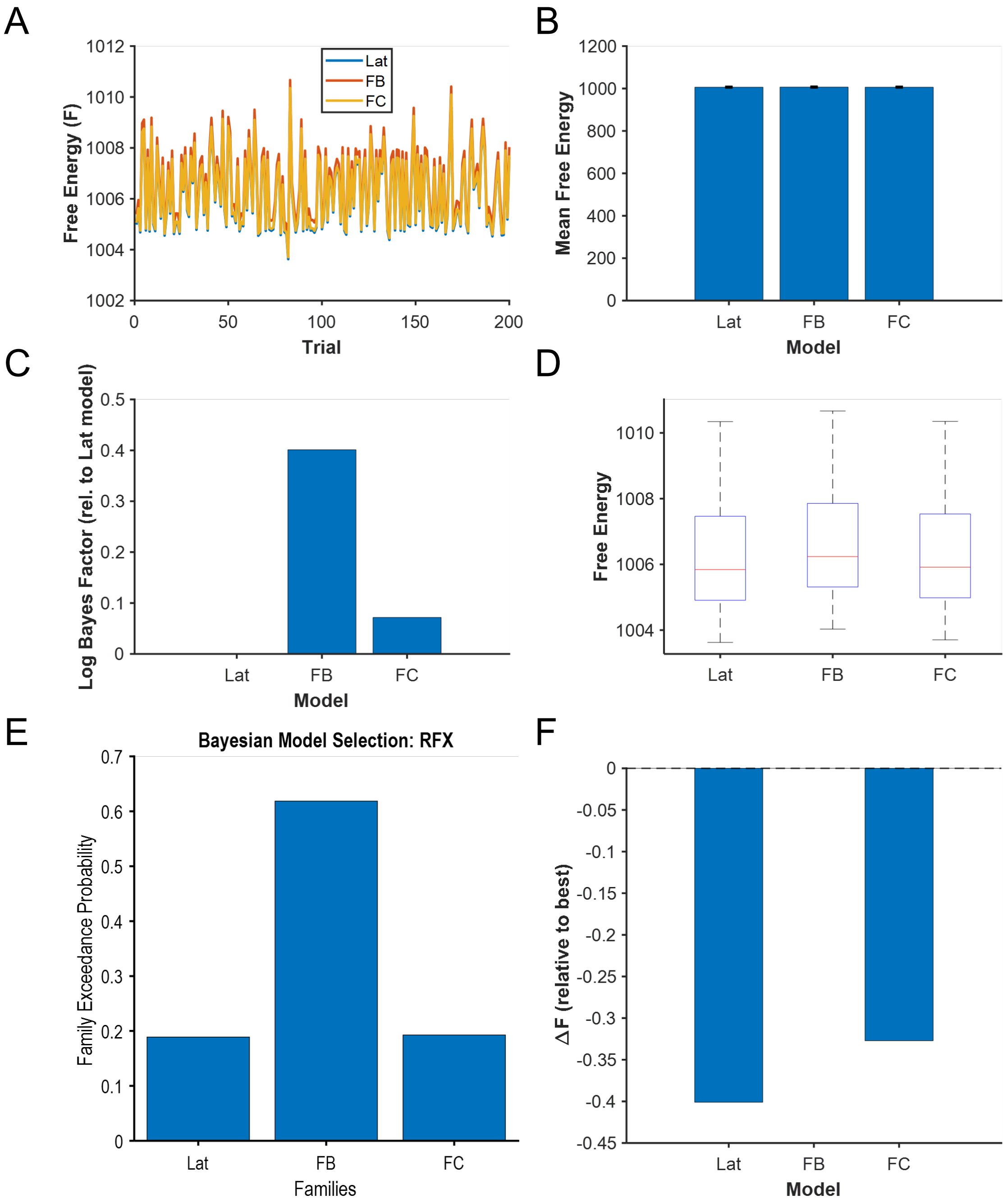


**Fig. S3. Results of DCM-CSD estimation in the period-doubling regime show forward-backward model as the winning model. (A)** Trial-wise free energy values across the three models: Lat (lateral), FB (forward-backward), and FC (fully connected). **(B)** Mean free energy per model with standard error. **(C)** Log Bayes factors computed relative to the Lat model, showing the highest evidence for the FB model. **(D)** Distribution of free energy across trials for each model (boxplot). **(E)** Bayesian model selection using random effects (RFX) indicating the FB model has the highest exceedance probability. **(F)** Within-subject parametric empirical Bayes (PEB) comparison shows that the FB model has the highest group-level free energy, with $\Delta F$ = -0.40 and -0.33 for Lat and FC models respectively, relative to FB. Together, these results show that CSD-based DCM failed to recover the ground-truth lateral connectivity model under period-doubling dynamics.

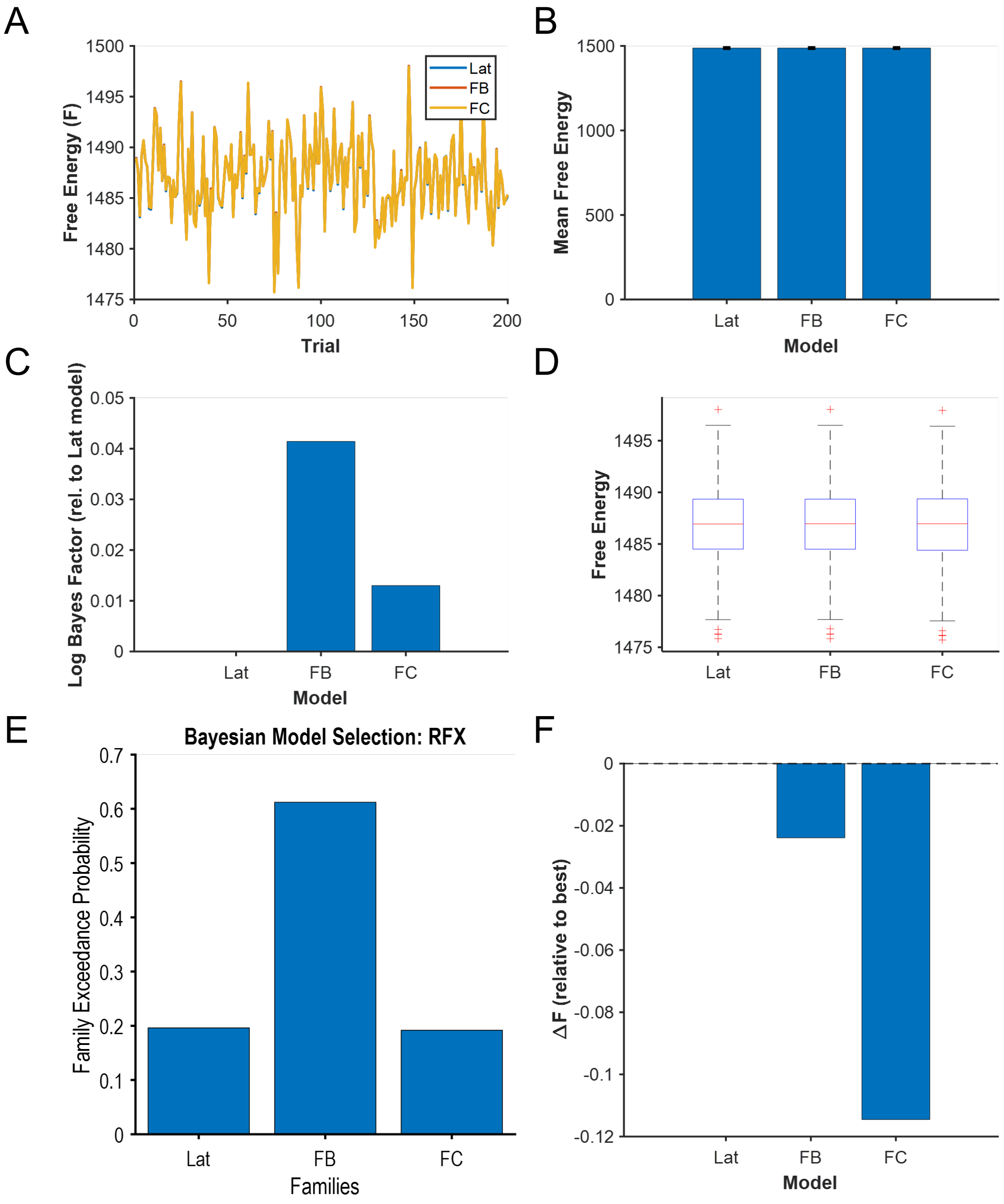


**Fig. S4. Results of DCM-CSD estimation in the chaotic regime. (A)** Trial-wise free energy values for each model: Lat (lateral), FB (forward-backward), and FC (fully connected). **(B)** Mean free energy across all trials with standard error. **(C)** Log Bayes factors relative to the Lat model indicate a slight preference for FB. **(D)** Boxplot of free energy distributions across trials. **(E)** Random-effects Bayesian model selection (RFX) identifies the FB model as the most likely, consistent with free energy metrics. **(F)** In contrast, within-subject PEB selects the Lat model as the winning model, with FB and FC showing ΔF = -0.02 and -0.11, respectively. These results suggest that while PEB can recover the ground-truth under chaotic dynamics, other inference methods may remain unreliable.

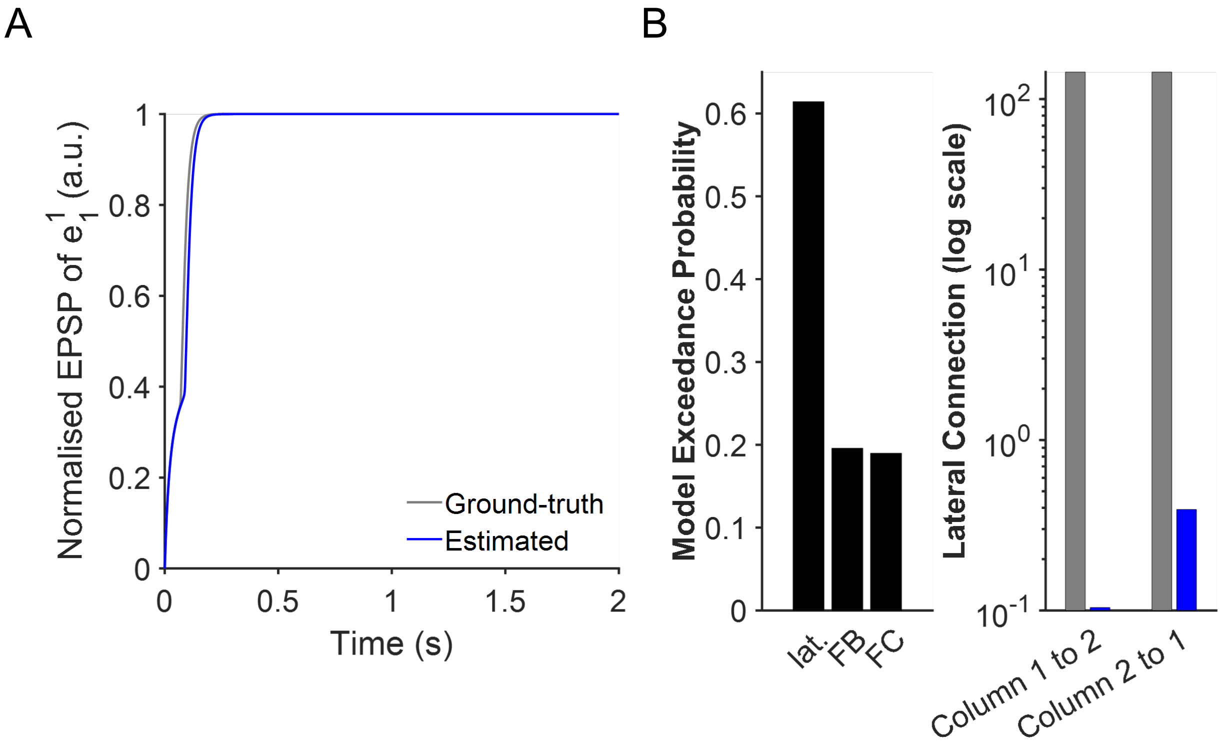


**Fig. S5. DCM accurately reproduces the single-steady-state output despite vastly different lateral connection strengths.** The thalamocortical model was configured to a single steady state by increasing self-excitation (*c7 = 3*) while maintaining balanced lateral couplings (E→E and E→I = 144). Simulations used noisy constant inputs of different amplitudes (Input 1 = 100.118 ± η, Input 2 = 0.112 ± η; *fs* = 1 kHz, *n* = 200 trials). **(A)** DCM-predicted normalised EPSP (blue) closely matches the ground truth (grey). **(B)** Random-effects BMS correctly identifies the lateral model (left). BMA-estimated lateral couplings (blue) are several orders smaller than the ground truth (grey, log scale), showing that near a fixed point, DCM reproduces normalised dynamics through compensatory scaling and SPM's default data normalisation rather than accurate absolute parameter recovery (right).

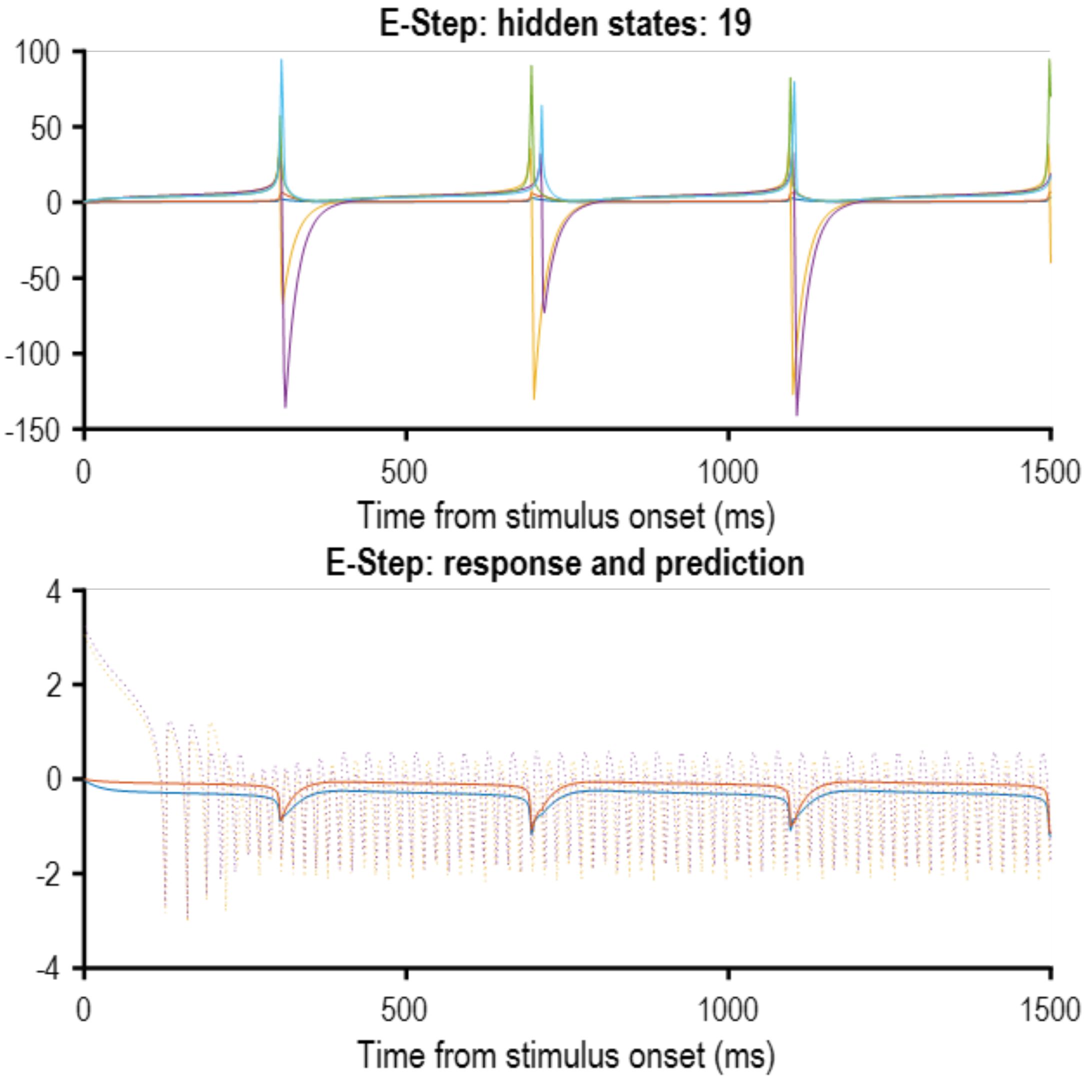


**Fig. S6. The effect of sampling frequency on dynamic causal modelling (DCM) parameter estimation in the period-doubling regime.** The synthetic neural signals were generated from the ground-truth cortical columnar model exhibiting period-doubling dynamics, sampled at a frequency of 500 Hz. Top: Inferred hidden states during DCM estimation, indicating the dynamics captured by the model. Bottom: Compares the original simulated response (dotted lines) with DCM's prediction (solid lines). Despite the SPM toolbox reaching algorithmic convergence, the predicted frequency failed to accurately follow the original period-doubling oscillations, highlighting a limitation of DCM in capturing precise neural dynamics with insufficient sampling frequencies.

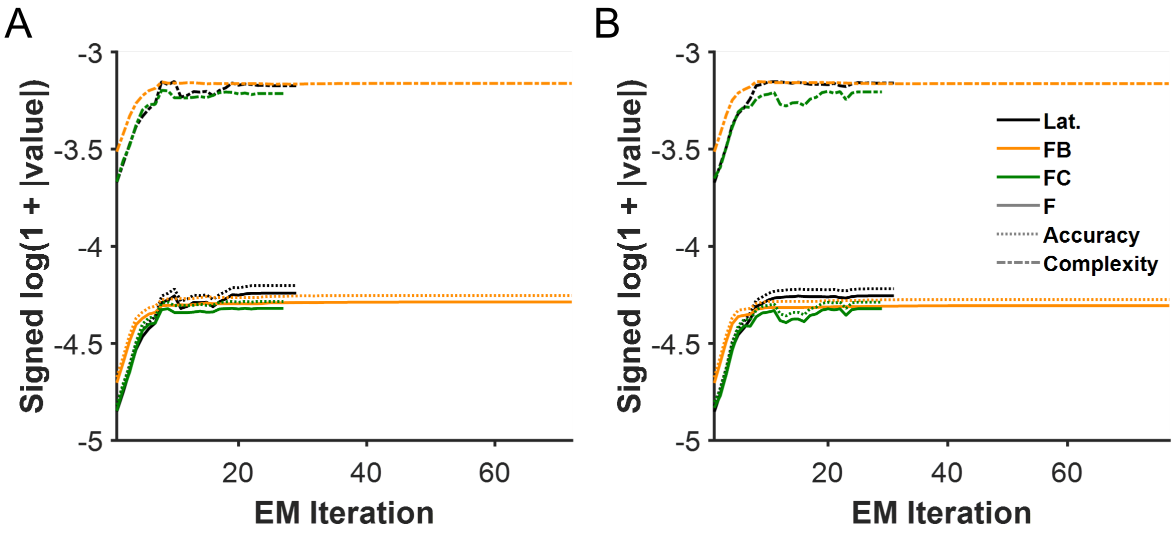


**Fig. S7. Tracking the evolution of variational free energy and its components during DCM estimation.** Free energy (solid lines), accuracy (dotted lines), and complexity (dash-dotted lines) are plotted across expectation maximation (EM) iterations for each model in the DCM model space: Lat. (black), FB (dark orange), and FC (dark green). **(A)** Period-doubling regime: the ground-truth (Lat.) model shows more irregular evolution in accuracy and free energy compared to other models. **(B)** Chaotic regime: Lat. And FB models display smoother convergence patterns. Despite convergence, model estimates fail to capture the ground-truth dynamics. This analysis highlights that free energy convergence alone may not guarantee accurate recovery of the temporal features of multistable dynamics.
